# Spike-dependent dynamic partitioning of the *Locus coeruleus* network through noradrenergic volume release in a simulation of nucleus core

**DOI:** 10.1101/2022.02.13.480256

**Authors:** Shristi Baral, Hassan Hosseini, Kaushik More, Thomaz M.C. Fabrin, Jochen Braun, Matthias Prigge

**Affiliations:** Research Group Neuromodulatory Networks, Leibniz Institute for Neurobiology, 39118 Magdeburg, Germany; Cognition Biology, Otto-von Guericke University, 39118 Magdeburg, Germany; Center for Behavioral Brain Sciences, 39118 Magdeburg, Germany

## Abstract

The *Locus coeruleus* (LC) modulates various neuronal circuits throughout the brain. Its unique architectural organization encompasses a net of axonal innervation that spans the entire brain; while its somatic core is highly compact. Recent research revealed an unexpected cellular input specificity within the nucleus that can give rise to various network states that either broadcast norepinephrine signals throughout the brain or pointedly modulate specific brain areas. Such adaptive input-output functions likely surpass our existing network models that build upon a given synaptic wiring configuration between neurons. As the distances between noradrenergic neurons in the core of the LC are unusually small, neighboring neurons could theoretically impact each other via volume transmission of NE. We therefore set out to investigate if such interaction could be mediated through noradrenergic alpha2-receptors in a spiking neuron model of the LC. We validate our model of LC neurons through comparison with experimental patch-clamp data and identify key variables that impact alpha2-mediated inhibition of neighboring LC neurons. Our simulation confirms a reliable autoinhibition of LC neurons after episodes of high neuronal activity that continue even after neuronal activity subsided. Also, dendro-somatic synapses inhibit spontaneous spiking in the somatic compartment of connected neurons in our model. We determined the exact position of hundreds of LC neurons in the mouse brain stem via a tissue clearing approach and, based on this, further determined that 25 percent of noradrenergic neurons have a neighboring LC neuron within less than a 25-micrometer radius. By modeling NE diffusion, we estimate that more than 15 percent of alpha2-adrenergic receptors fraction can bind NE within such diffusion radius. Our spiking neuron model of LC neurons predicts that repeated or long-lasting episodes of high neuronal activity induce partitioning of the gross LC network, and reduces the spike rate in neighboring neurons at distances smaller than 25 micrometers.

As these volume-mediate neighboring effects are challenging to test with the current methodology, our findings can guide future experimental approaches to test this phenomenon and its physiological consequences.

## Introduction

The *Locus coeruleus* (LC) is a brain stem nucleus that is the primary source of norepinephrine (NE) in the central nervous system. It comprises a unique neuronal architecture with an extensive and diffuse axonal innervation throughout the entire brain and spinal cord, while the somatic core consisting of several hundred neurons is highly compact (Szabadi 2013). Research on the noradrenergic system has revealed astonishing insights on how a single neuromodulator such as NE can pointedly impact diverse brain functions ranging from arousal to pain sensation to memory encoding and sensory processing (Carter et al. 2010; Hirschberg et al. 2017; Takeuchi et al. 2016; Martins and Froemke 2015). In addition, much research is focused on understanding how expression profiles of adrenergic receptors in their respective axonal terminal fields are correlated to mental disorders such as attention deficit, clinical depression, or drug abuse; reviewed in (Poe et al. 2020). Distinct catecholaminergic receptor subtypes can be activated through a dynamic modulation of an extracellular NE gradient in the vicinity of a synaptic release site (Takeuchi et al. 2016; Zhang et al. 2013). In this respect, the type of NE release site in axonal terminals has been controversially discussed in the past; to one extrem a pure volume transmission was assume as main mode of action, and on the other extrem a purely wiring-like transmission (Descarries, Watkins, and Lapierre 1977; Latsari et al. 2002; Olschowka et al. 1981; Pickel et al. 1981; Nirenberg et al. 1996).

On the opposite, less research has focused on the synaptic wiring modes within the LC core itself. Primarily, such research interest was hampered by the canonical view that LC activity is mainly based on autonomous pacemaking of a homogenous noradrenergic cell population. Yet, recent technological advancements in single-cell genetics, viral tracing, and optical manipulation of neurons begin to challenge this view. Already a surprising heterogeneity in the LC cell population regarding their projection-specificity and physiological function has been described (Chandler, Gao, and Waterhouse 2014; Uematsu et al. 2017; Hirschberg et al. 2017). Nevertheless, we are still missing a framework on how individual LC neurons are computing their brain-wide inputs to the nucleus and how this information is relayed within the nucleus and thereby to the entire brain (Poe et al. 2020). As the somas in the nucleus are in unusually close proximity to each other, volume-mediated interactions are perceivable. Previous work on catecholaminergic neurons in the substantia nigra and ventral tegmental area argue for such volume interaction between neurons as a key mechanism to fully describe neuronal activities in these areas (Rice and Cragg 2008; Rice and Patel 2015; Cragg and Rice 2004). Early work in LC focused primarily on electronic coupling via gap junction as the main mode of connection (Ishimatsu and Williams 1996). Gap junctions are most prominent in the dendritic compartment and decrease in abundance with aging (Christie and Jelinek 1993; Christie, Williams, and North 1989). They give rise to subthreshold oscillation that can synchronize the entire nucleus during postnatal periods (Ishimatsu and Williams 1996). Yet, recent high-density silicon probes recordings confirm that at older age neurons from different subsets of the LC exhibit asynchronous activity (Totah, Logothetis, and Eschenko 2020). Here, correlated activity patterns are observed within these subsets of LC neurons at different timescales: from sub-milliseconds to milliseconds to periodic oscillation in the range of seconds. Such a wide range of temporal interactions between LC neurons is beyond a pure gap junction coupled neuronal network.

Indeed, several neuromodulators such as galanin, serotonin, neuropeptides and NE itself are known to also impact LC network activity on the timescale of seconds and even minutes (Tillage et al. 2020; Bai et al. 2018; Segal 1979; Aghajanian, Cedarbaum, and Wang 1977). Electrophysiological studies clearly demonstrated a strong NE-mediated control of the LC network via high-affinity alpha2-receptor (Singewald 1998; Singewald et al. 1994; Lakhlani et al. 1997; Wagner-Altendorf, Fischer, and Roeper 2019). Infusion of antagonists leads to hyperpolarization that is mediated through a NE-sensitive potassium conductance (Swanson 1976; Lee, Rosin, and Bockstaele 1998; Torrecilla et al. 2013). This decrease in excitability is mediated via G_i/o_ pathways that trigger the G-protein coupled inward rectifying potassium channel GIRK2 (also called Kirk3) thereby controlling afferent activation of LC neurons as well as the network state of the entire nucleus (Travagli, Dunwiddie, and Williams 1995; Travagli, Wessendorf, and Williams 1996; Williams, North, and Tokimasa 1988; Aghajanian, Cedarbaum, and Wang 1977; Lüscher and Slesinger 2010). So far, these questions have been solely approached through a drug-based experimental design, where a drug effect is either studied with conventional patch-clamp recordings on a single cell level or at the other extrem, on the level of the entire LC network. As our view of the LC network shifts to a microcircuitry level of functional specific subnetworks within the LC, spiking patterns of individual neurons in their respective subnetworks become more critical for our understanding. Episodes of high spiking rate (burst-like activity) in individual neurons can evoke elevated intracellular calcium levels in soma that surpass the baseline level seen during tonic activity (Huang et al. 2012). These high calcium levels can trigger the release of NE-containing vesicles from dendritic and somatic compartments (Huang et al. 2007; 2012). It is challenging to evaluate how synaptic and extracellular NE levels influence the LC network and neighboring neurons with the current technologies. We, therefore, set out to employ a Hodgkin-Huxley-based model to study NE-mediated local effects with the nucleus.

## Methods

### Acute brain slice preparation

All procedures were approved by the Landesverwaltungsamt / Halle, and conformed to German regulation. All studies were performed in compliance to animal protocol 42502-2-1545 LIN. Acute brain slices for electrophysiology where obtained from Ai9 x DbH-Cre mice (Jax# 007905 and 033951) on a C57Bl6/J background. For tissue clearing mice from an Ai9 x PV-Cre were used (Jax# 007905 and 17320).

Mice were injected intraperitoneally with pentobarbital (Narcoren^®^, Boehringer Ingelheim Vetmedica GmbH, D-55216 Ingelheim, Germany,130 mg/kg, i.p.) and perfused with carbonated (95% O2, 5% CO2) ice-cold slicing solution (in mM: 2.5 KCl, 11 glucose, 234 sucrose, 26 NaHCO3, 1.25 NaH2PO4, 10 MgSO4, 2 CaCl2; 340 mOsm). After decapitation, 300-μm coronal brain stem slices were prepared in carbonated ice-cold slicing solution using a vibratome (Leica VT 1200S) and allowed to recover for 20 min at 33 °C in carbonated high-osmolarity artificial cerebrospinal fluid (high-Osm ACSF; in mM: 3.2 KCl, 11.8 glucose, 132 NaCl, 27.9 NaHCO3, 1.34 NaH2PO4, 1.07 MgCl2, 2.14 CaCl2; 320 mOsm) followed by 40 min incubation at 33 °C in carbonated ACSF (in mM: 3 KCl, 11 glucose, 123 NaCl, 26 NaHCO3, 1.25 NaH2PO4, 1 MgCl2, 2 CaCl2; 300 mOsm). Subsequently, slices were kept at RT in carbonated ACSF until use. The recording chamber was perfused with carbonated ACSF at a rate of 2 ml min–1 and maintained at 32 °C.

### Brain slice electrophysiology

Whole-cell patch-clamp recordings were performed under visual control using oblique illumination on a Slicecope Pro2000 (Scientifica) equipped with a 12-bit monochrome CMOS camera (Hamamatsu Model OrcaFlash). Borosilicate glass pipettes (Sutter Instrument BF100-58-10) with resistances ranging from 3–7 MΩ were pulled using a laser micropipette puller (Sutter Instrument Model P-2000). Pipettes were filled using standard intracellular solution (in mM: 135 potassium gluconate, 4 KCl, 2 NaCl, 10 HEPES, 4 EGTA, 4 Mg-ATP, 0.3 Na-GTP; 280 mOsm kg–1; pH adjusted to 7.3 with KOH) Whole-cell voltage-clamp recordings were performed using a MultiClamp 700B amplifier, filtered at 8 kHz and digitized at 20 kHz using a Digidata 1550A digitizer (Molecular Devices).

### HEK cells electrophysiology

Human embryonic kidney (HEK) cells stably expressing G-protein rectifying potassium channels (GIRK 1/2 subunits), kindly provided by Dr. A. Tinker UCL London, GB, and Dr. S. Herlitze were maintained at 37 °C in Dulbecco’s modified Eagle’s medium, 4.5 g/l D-glucose, supplemented with 10% fetal bovine serum (Gibco) and penicillin/streptomycin incubated under 5% CO2. Cells were cultured in a 24-well plate, on 12 mm glass coverslips.

Whole-cell patch-clamp recordings were performed under visual control using a Slicecope Pro2000 (Scientifica) equipped with a 12-bit monochrome CMOS camera (Hamamatsu Model OrcaFlash). Borosilicate glass pipettes (Sutter Instrument BF100-58-10, 2-5 MΩ) were pulled using a laser micropipette puller (Sutter Instrument Model P-2000). Pipettes were filled using the internal solution as follows: 100 mM potassium aspartate, 40 mM KCl, 5 mM MgATP, 10mM HEPES-KOH, 5 mM NaCl, 2 mM EGTA, 2 mM MgCl2, 0.01 mM GTP, pH 7.3 (KOH). The bath was filled using the following external solution: 20 mM NaCl, 120mM KCl, 2 mM CaCl2,1mM MgCl2, 10 mM HEPES-KOH, pH 7.3 (KOH). Whole-cell voltage-clamp recordings were performed using a MultiClamp 700B amplifier, filtered at 8 kHz and digitized at 20 kHz using a Digidata 1550A digitizer (Molecular Devices). The following protocol was used during the IV curve recordings: fifteen sweeps starting from −80 mV (Δ+10 mV). The cells were clamped on a holding potential at −60 mV.

### Immunohistochemistry

Mice were transcardially first perfused with phosphate-buffered saline (PBS 1x) and then with 4% paraformaldehyde. The brains were extracted and stored overnight in 4% PFA followed by soaking in 30% sucrose to avoid forming water-crystals during slice sectioning at 4°C. Brains were sliced on a microtome (LEICA SM2010R) to 40 μm thick coronal sections. Afterward, brain slices were first permeabilized and blocked by incubating in PBST (Phosphate-buffered saline with 0.3% Triton-X) containing 10% horse serum for 2 hours at room temperature to prevent non-specific binding of antibodies. Sections were incubated in a solution containing the primary antibodies, 2% horse serum, and 0.3% Triton-X for 24hrs at 4°C. The primary antibodies used were guinea pig anti-TH (Synaptic Systems, Catalog No. 213104) with 1:500 dilution and rabbit anti-DBH (ImmunoStar, Catalog No. 22806) with 1:1000 dilution. Following three washes in 1x PBS for 10 min the sections were then incubated in a solution containing Cy3 anti-guinea pig IgG, (Jackson ImmunoResearch, Catalog No. 706-165-148), Cy5 AffiniPure Donkey Anti-Rabbit IgG, (Jackson ImmunoResearch, Catalog No. 711-175-152) both in 1:200, 5% horse serum and 0.05% Triton-X for 2 hours at room temperature. The sections were then washed thrice with 1xPBS for 10 min and then incubated in a solution containing DAPI at a dilution of 1:30000. All the above-mentioned solutions were made in 1x PBS. The sections were mounted on glass slides and coverslips containing DPX mounting media. Images were acquired on a Leica SP8 confocal microscope using 63x oil-immersion objectives (NA 1.4). Images were analyzed using ImageJ (https://imagej.nih.gov/ij/). For images in figure 1, we overlay the reference atlas form *The Mouse Brain in Stereotaxic Coordinates 3rd Edition Franklin & Paxinos* at anterior-posterior axis −5.40mm.

**Figure 1.**
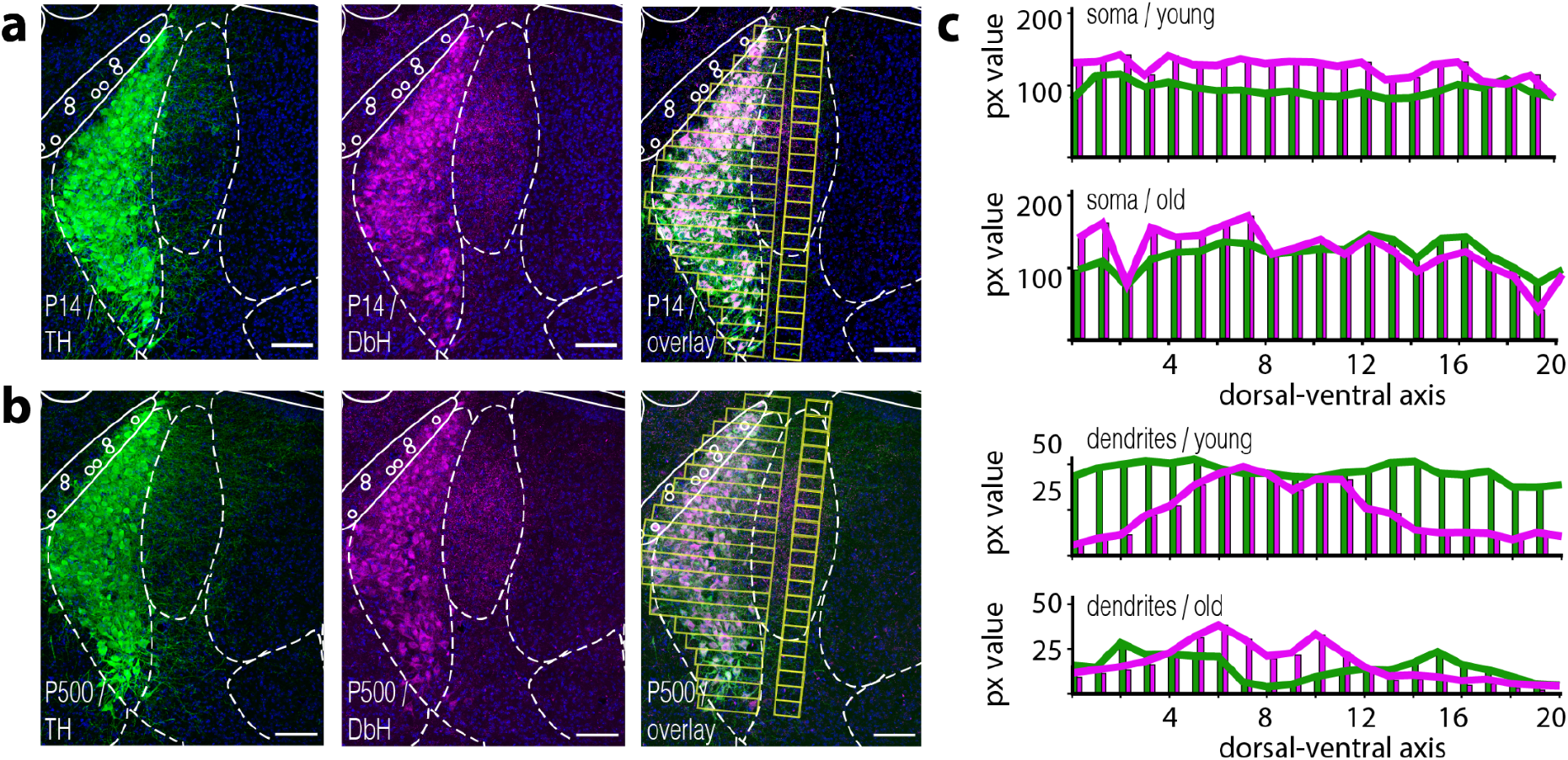
Tyrosine hydroxylase and Dopamine-beta-Hydroxylase expression largely overlap in somatic and dendritic compartments in the Locus coeruleus. (**a**) Shows immunostaining against key enzymes in the synthesis pathway of NE. White dotted lines show an overlay mask of the corresponding section from The Mouse Brain in Stereotaxic Coordinates 3rd Edition Franklin & Paxinos **(a)** Confocal image of immunofluorescence against tyrosine hydroxylase (TH / g*reen*) from a brain stem slice of a young P14 mouse (anterior-posterior axis: −5.40 mm). The fluorescence signal is present in the soma as well as in the dendritic arbor (here more dominantly in the medial part); We observed a similar spatial distribution for the fluorescence signal from immunostainings against the dopaminebeta-hydroxylase (*magenta*). **(b)** depict the same immunostainings as in (a) on brain slices obtained from an old P500 mice. A similar spatial expression profile for both enzymes is visible when compared to slices from a young P14 mouse (a). **(c)** Shows a quantification of average pixel (px) intensities from the *yellow-colored ROIs* displayed in TH/DbH overlay image in a and b. ROIs are placed above the somatic or dendritic compartments, respectively. **(c)** Shows average intensity of pixel values from ROIs along the dorsal-ventral axis for soma or dendrites. Scale bar 100μm

### Tissue-clearing

Sectioning of 4% PFA fixated mouse brains were performed in a coronal configuration using a brain mold to obtain 2-mm thick slices that contain the major part of the LC nucleus (AP 4.4 to 6.4mm) (brain matrix, 3D printed). Brain slices were then washed in PBS for 1h and incubated at different series of mixtures prepared from tetrahydrofuran (THF, Sigma-Aldrich, Catalog No.186562-1L):dH2O + 2% Tween-20 to dehydrate brain slices. The pH of the THF solution was adjusted to 9.0 by adding triethylamine. The slices were washed in 50% THF solution for 4h at 4°C followed by 70%, 90%, and 100% THF solutions under the same conditions. To ensure complete dehydration, we washed the section with 100% THF twice. Slices were then bleached in chilled 5% H_2_O_2_ in THF overnight at 4°C. The next day, the slices were rehydrated in the reverse order by washing them in 100%, 90%, 70%, and 50% THF solutions as above for 4h at 4°C. Following this, slices were washed twice in a PBS / 0.2% TritonX for 1h each wash. Slices were then permeabilized in PBS / 0.2% TritonX / 20% DMSO / 0.3 M glycine at 4°C for 2 days and then blocked in PBS/0.2% TritonX / 10% DMSO / 6% horse serum at 4°C for another 2 days. Samples were kept in primary antibody (guinea pig anti-TH Synaptic Systems, Catalog No. 213104) dilution in PBS / 0.2% Tween-20 / 5% DMSO / 3% horse serum with 10 μg/ml heparin for two days at 4°C. Primary antibodies were diluted at 1:400. Following this treatment, samples were washed in PBS / 0.2% Tween-20 with 10 μg/ml heparin for 1h at 4°C for 4x and then kept in secondary antibody (Cy3 anti-guinea pig IgG, Jackson Immuno Research, Catalog No. 706-165-148) in 1:150 dilution in PBS / 0.2% Tween-20 / 3% horse serum with 10 μg/ml heparin at 4°C for 2 days and washed in PBS / 0.2% Tween-20 with 10 μg/ml heparin for 1h at 4°C for 4x. For imaging, samples were embedded in 0.8% phytagel and dehydrated in the THF/dH20 series exactly as described before. After the dehydration, the samples were transferred to Ethyl Cinnamate (ECi) at room temperature for clearing. ECi was replaced with fresh ECi the following day, and samples were stored at room temperature.

### Light-sheet microscopy

The light-sheet microscopy was performed using LaVision Biotech Ultramicroscope II (Miltenyi Biotec, Bergisch-Gladbach, Germany) which was equipped with a Zyla 4.2 PLUS sCMOS camera (Oxford Instruments, Abingdon-on-Thames, UK). The embedded brain slice was placed in the sample chamber containing Ethyl Cinnamate (ECi). The brain volume was sampled through a 4x objective (LaVision BioTec LVMI-Flour 4x/0.3). The objective was configured to match the refractive index of ECi (1.558), for optimal imaging. For excitation, we used light from a EXW-12 extreme laser (NKT Photonics, Birkerød, Denmark) which was guided through the appropriate excitation filters (AHF Analysentechnik AG, Tübingen, Germany) for Cy3 fluorescence. We started acquiring the image using ImSpector software (Miltenyi Biotec, Bergisch-Gladbach, Germany) after setting the parameters such as sheet width, z-step sizes, number of tiles and zoom. For post-processing and segmentation we used Imaris software (version 9.72 Bitplane, Zurich, Switzerland).

### Modeling

Based on cortical membrane kinetics, Alvarez et al. 2002 proposed a two-compartment mathematical model of LC cells, with soma and dendrite connected via a gap junction on the dendrite. We employed this model as an initial starting point and extended the model to incorporate quantal release of NE, which eventually leads to the opening of GIRK channels and inhibition of the neighboring cell and/ or itself via alpha2-receptor located in the somatic area. The release event is coupled with the amount of cytosolic calcium [Ca^2+^] in the cell.

### Dynamic equation of state variables

Voltage change in the soma is driven by the balance of currents, including electrode current *I_elec_*, HH-sodium current *I_Na_*, HH-potassium current I_K_, persistent sodium current I_P_, Ca^2+^-driven potassium current I_AHP_, voltage-driven calcium current I_Ca_, synaptic current from GIRK conductance I_GIRK_, current from dendrite I_dend_, leak current I_L_, and independent random current η(t).

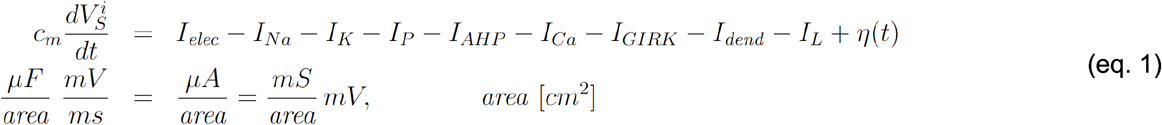

Voltage change in the dendrites are driven by current from soma I_soma_, gap junctions from other dendrites I_GJ_, and dendritic leak current I_LD_

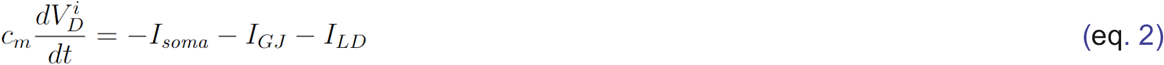

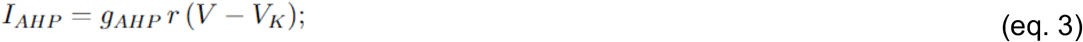

The Ca^2+^ concentration changes over time as a function of voltage.

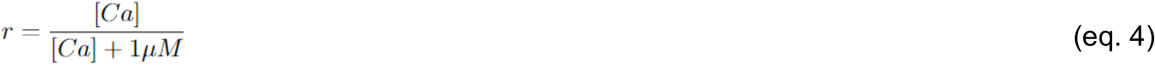

Specifically, membrane depolarization during a spike allows Ca^2+^ to enter the neuron. As described above, the opening and closing of the gAHP potassium conductance shadow the cytosolic Ca^2+^ concentration.

The steady-state of the calcium is

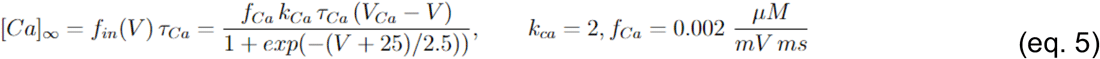

And the update rule is:

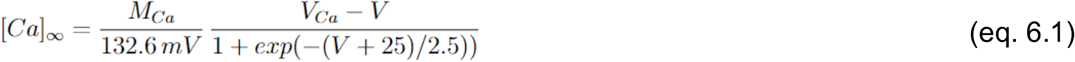

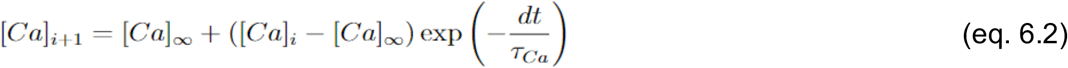

As the voltage-dependent factor in the expression of [Ca]∞ peaks at 132.6mV for V_Ca_ = 120mV, it is convenient to define a new parameter, M_ca_ = 42.4 μM and *τ_Ca_*= 80ms (supplemental figure 1)

### Dendro-somatic and soma-somatic NE release and alpha2-receptor activation, and GIRK conductance

Action potentials in the presynaptic neuron results in NE release of dense core vesicles from presynaptic sites, which are found at the soma and at nearby dendrites. The release fraction R_rel_ depends on the level of presynaptic calcium at the time of the presynaptic action potential.

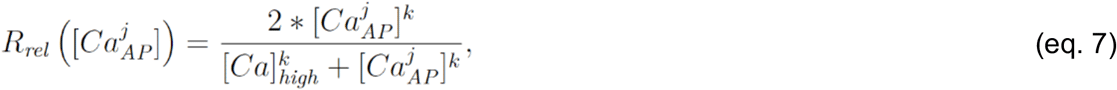

[Ca^j^AP] is presynaptic cytosolic calcium at the time of the action potential. [Ca]_high_=1.3 μM is the amount of cytosolic calcium accumulated when the cell fires at 20 Hz. The exponent, k=1, modulates the release.

Once released, NE binds to alpha-2 receptors in the postsynaptic membrane, which through G-protein signaling opens an inwardly-rectifying potassium channel (GIRK: G-protein-coupled inward-rectifying potassium channel). The effective time course of the opening and closing of this synaptic conductance is comparatively slow (Williams, Bobker, and Harris 1991). We approximate this time-course with a double-exponential function with *τ*_fest_= 300 ms and *τ*_slow_= 350 ms as well as measure the voltage-dependence of GIRK current in a HEK cells line that is stably expressing GIRK1/2 subunit channel (supplemental figure 2) (Williams, Bobker, and Harris 1991; Eickelbeck et al. 2020).

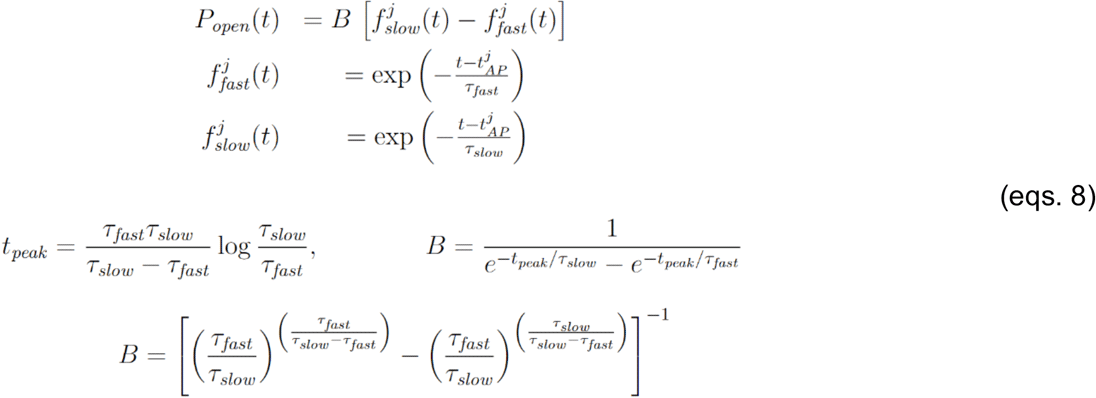

The absolute value G_rect_ of this rectifying conductance is not constant, but changes with the postsynaptic membrane potential. Accordingly, we need to consider this factor in the equation.

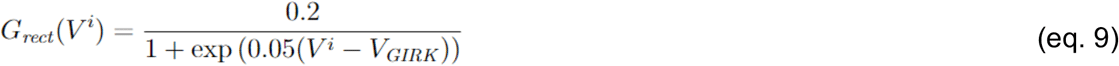

G_rect_ is given in μS per mm^2.^

Combining all three factors for multiple successive presynaptic action potentials AP_1_, AP_2_,…, the GIRK conductance is given then by:

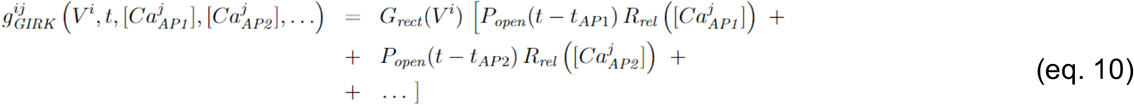

Release from or near soma and binding near soma. This hyperpolarizes the membrane potential of the neighboring soma. S_ij_ is the strength of the synaptic connection from neuron j to neuron i.

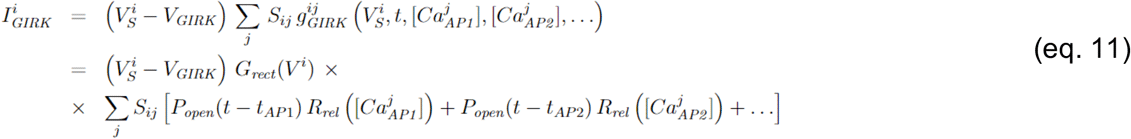

### NE diffusion

To estimate the effect of NE released into the extracellular space, we computed the time-course of NE concentration at some distance from a release site as by:

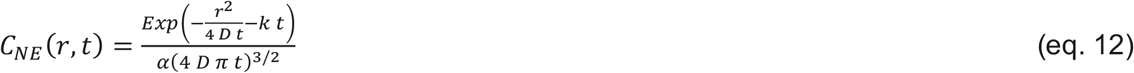

where r is distance, t is time, D=3.4e-6 cm^2/s is the diffusion constant, and k = 20 Hz is the reuptake constant (Rice and Nicholson 1995). The fraction of NE-binding alpha2-receptors was computed from

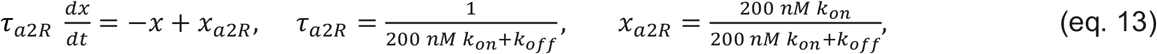

where k_on_ = 1.0 and k_off_ = 1.25 nM are the kinetic constants for NE binding and unbinding. The resulting time course and voltage dependence of GIRK conductance was fitted with a double-exponential function with *τ*_fest_= 1400 ms and *τ*_slow_= 1450 ms and an attenuation factor of 0.07 (supplemental figure 2).

To take into account the kinetics of transmitter binding and release, the resulting time-course was convolved with a low-pass filter (‘alpha function’) 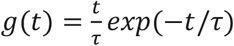 with characteristic time *τ* = 50ms.

### Data analysis

All data were analyzed in Python. For extraction of action potential features (simulated and experimental), we used the Allen Institute ephys package. The generalized leaky integrate and fire models (GLIF) were applied. Additional analyses were performed using custom Python scripts, all of which are available on the laboratory GitHub account.

## Results

We first set out to confirm that LC neurons express both key enzymes to catalyze NE in somatic and dendritic compartments throughout their lifetime. Immunostaining against tyrosine hydroxylase (TH), as well as dopamine-beta-hydroxylase (DbH) in young mice (P14) and old mice (older than P500), revealed expression of both enzymes in all cells at both ages (figure 1 a, b). Both stainings also highlight the compact nature of the LC nucleus itself. Furthermore, a dense dendritic arbor directed towards the medial-rostral axis becomes visible (Ishimatsu and Williams 1996). Image overlay of the fluorescence channel from TH and DbH confirms an overall co-expression of both enzymes throughout the somatic compartment within the LC nucleus (see figure 1 a and b overlay). Surprisingly, we observed a differential spatial expression levels of TH and DbH in the dendritic arbor. Here, both poles of the LC (dorsal and ventral) appear to have protruding dendrites that mainly display a TH immunosignal. The central part of the nucleus seems to extend dendrites that preferentially express DbH (figure 1 c, 3 animals). The dendritic expression profile for the DbH enzymes was also similar for young and old mice along the anterior-posterior axis (supplemental figure 3)

As it is difficult to experimentally approach somatic or dendritic NE release from a single LC neuron and investigate how it affects spike activity in neighboring LC neurons, we decided to develop a computational model that allows us to study three modes of NE-mediated interactions. The first mode was via dendro-somatic synapse (Nirenberg et al. 1996); secondly, we considered a somatic autoinhibition (Andrade and Aghajanian 1984; Wagner-Altendorf, Fischer, and Roeper 2019), and thirdly we investigated if a NE volume diffusion from a somatic release point can lead to an inhibition of a neighboring neuron.

To realistically model the distribution of the distances between LC neurons, we employed a tissue clearing approach to map the positions of noradrenergic neurons in the LC. First, we prepared 2-mm thick brain stem slices from PV-Cre x Ai9 animals. We then used a modified iDISCO protocol to clear the entire brain stem slices and image the slice volume via fluorescence light sheet microscopy (see methods for details). After acquiring images of the entire brain stem, we visualize anti-TH+ and PV-dTomato neurons with high contrast (figure 2 a). We segmented the LC and resolved positions of 220 to 439 anti-TH+ neurons (4 nuclei, 2 animals) (figure 2 b). Based on the x-y-z coordinates of individual anti-TH+ neurons, we compute the nearest-neighbor (NN) distances for each cell in all four analyzed nuclei. The distribution of the NN-distances peaks at distances of beta-hydroxylase around 25μm (median 41.34μm) (see figure 2 c).

**Figure 2.**
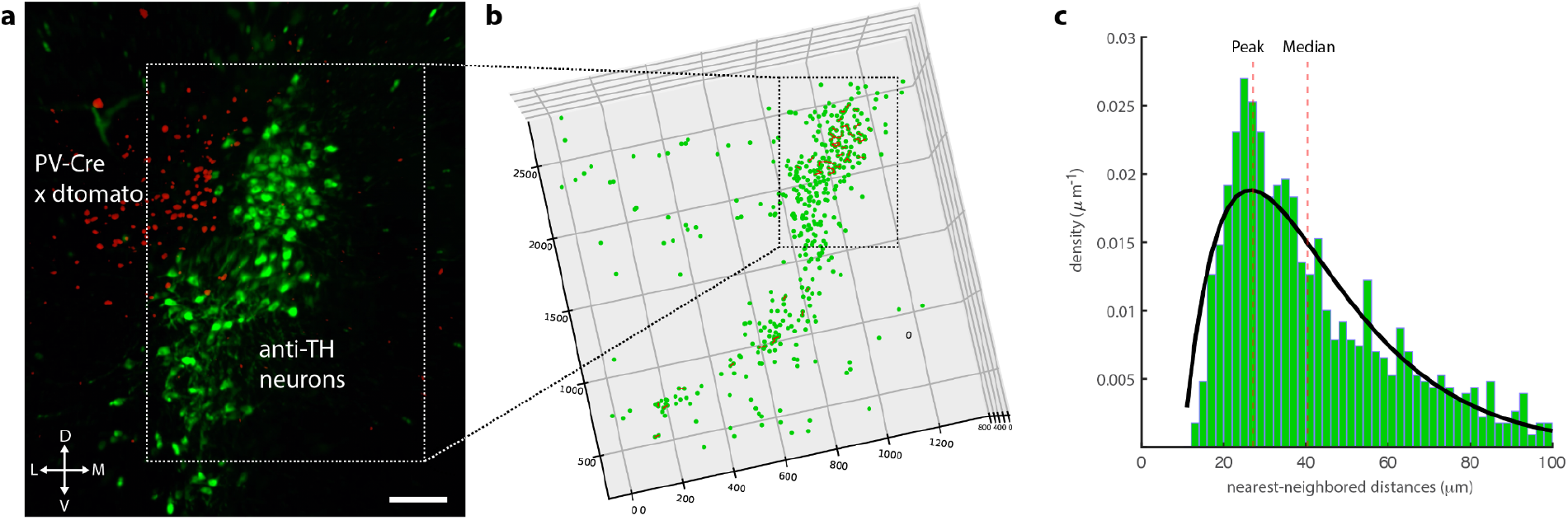
The largest fraction of noradrenergic neurons have neighboring LC neurons at a distance of 25 micrometers. **(a)** A typical fluorescent volume from a 2-mm cleared brain stem slice. Red dtomato fluorescence comes for parvalbumin-positive neurons while pixel intensities mapped in green at anti-TH / CY3 immunopositive neurons (scale bar 100μm). **(b)** Cell segmentation was manually performed with IMARIS software and validated through two observers. The mean number of recovered LC neurons was 289±45 (n = 4, 2 animals). Anti-TH positive neuron had a mean sphericity of 0.83±0.12 and a mean volume of 4971±.233μm^3^. **(c)** Nearest-neighbor distance histogram of pooled data sets from all 4 LC nuclei. Distances between neurons with less than 25μm are connected through a red line. Nearest-neighbor distribution was fitted to a double exponential function (γ = 1.38, λ_long_=16.84 and λ_short_ =16.23). The model predicts that the highest number of neurons are in bins between 24 to 28μm. The peak of NN distribution and median are marked as dotted lines.

To investigate the possible effects of a noradrenergic interaction between LC neurons on a single cell level, we choose as a starting point a previously published mathematical model that describes the electronic coupling between two LC neurons via gap junctions (Alvarez et al. 2002). This model recapitulates frequency-dependent synchronization between LC neurons as an electrotonic coupling strength between dendrites. Here, strong gap-junction coupling between dendritic compartments, as described experimentally in young mice (Christie and Jelinek 1993; Christie, Williams, and North 1989; Ishimatsu and Williams 1996), lead to synchronization of spontaneous spiking for all frequencies; while weak dendritic coupling promotes spike synchrony only at higher spiking frequencies. Comparison of our initial model under a weak electrotonic coupling regime reiterates our current-clamp recordings from noradrenergic neurons from acute brain slices prepared from mice older than P70. The spontaneous spike rate at resting potential was 4.2±1.77Hz (n = 6 neurons, 2 animals) while our model estimates 4.3 ± 0.03 Hz (figure 3 a). Overall features of action potentials between our model and experimental data largely coincide (figure 3 b). To emulate different excitability levels of LC neurons, we recorded the spike rate during a three hundred millisecond long current pulse at various amplitudes (figure 3 c, *upper trace*). Also, in our model, current steps evoked an increased spike rate (figure 3 c, *lower trace*). As expected, spike rates increased with amplitudes of injected current. Quantification of spike rate at various current step amplitudes revealed an overall linear dependence in the frequency-current relation (FI-curve) for experimental and simulated data (compare figure 3 d1 and d2). Interestingly, we observed a higher spike rate for the two initial spikes and an overall steeper slope in the FI-curve for these early spikes compared to the following spikes (supplemental figure 3 a). We also found such an increased spike rate for the initial spikes in our model; this effect gets more pronounced at higher current amplitudes for experimental and simulated data (compare supplemental figures 4 a and b).

**Figure 3.**
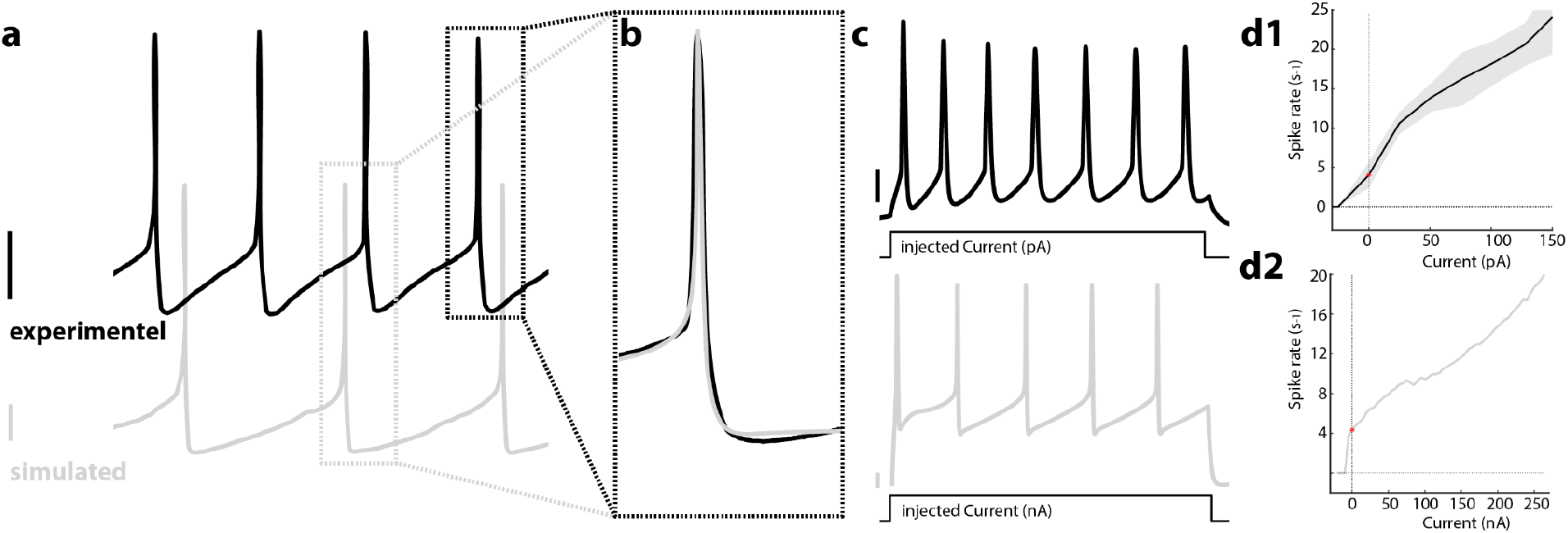
LC neuron model recapitulates essential spontaneous spiking, action potential characteristics and current-evoked frequency dependency. **(a)** Spontaneous neuronal spiking in an LC neuron recorded in an acute brain slice from P70 mice via whole-cell patch-clamp *(upper black trace*). Action potentials show typical long after-hyperpolarization (AHP) that give rise to a 4.2 Hz spontaneous spiking rate. Trace below shows simulated LC neuron that exhibit similar spontaneous spiking rate and slow AHP. **(b)** Overlay of experimental and simulated action potentials exhibit a similar temporal profile. **(c)** The current-evoked increase in membrane potential evokes a barrage of action potentials in a recorded and simulated neuron (300ms current injections). **(d1)** Illustrates the spike rate of recorded LC neurons under different current step amplitudes (error margin as SEM; n = 12 cells, 2 animals); **(d2)** Simulated LC neurons exhibit an increase in spike rate for different amounts of injected current. Note the different current units in d1 and d2 due to the different cell sizes of experimental and simulated neurons. Scale bars 10mV for experimental neurons and 20mV for simulated neurons.

To simulate a noradrenergic modulation between LC neurons, we extended our model with a calcium-dependent release of NE. We considered a NE release site as a point source of NE secretion that either mimics a dendro-somatic synapse or a somatic release site that can either retroact on the secreting neuron (self-inhibition) or effect neighboring neurons via diffusion (neighboring inhibition). High spiking rates lead to membrane fusion of NE-containing large-dense vesicles at the soma (Huang et al. 2007; 2012). The voltage-dependent rise in calcium is modeled by equations 3, 4, 5 and 6 (see Methods). Based on amperometric recordings in acute brain slices, we assumed an initial extracellular NE concentration after vesicle fusion of 200nM (Huang et al. 2012). We combined the time courses of several molecular events in equation 10 i.e. binding of NE to alpha2-receptors, initiation of G_i/o_ signaling cascade, and the subsequent release of G_βγ_ dimer that increases GIRK channel conductance. The dependence of GIRK conductance and intracellular calcium levels is shown in figure 4a. For a given calcium level (dotted line in figure 4a) the temporal development of GIRK open probability (P_open_) is modeled based on electrophysiological characterization of GIRK current in LC neurons (supplemental figure 2) (Williams, Bobker, and Harris 1991).

**Figure 4.**
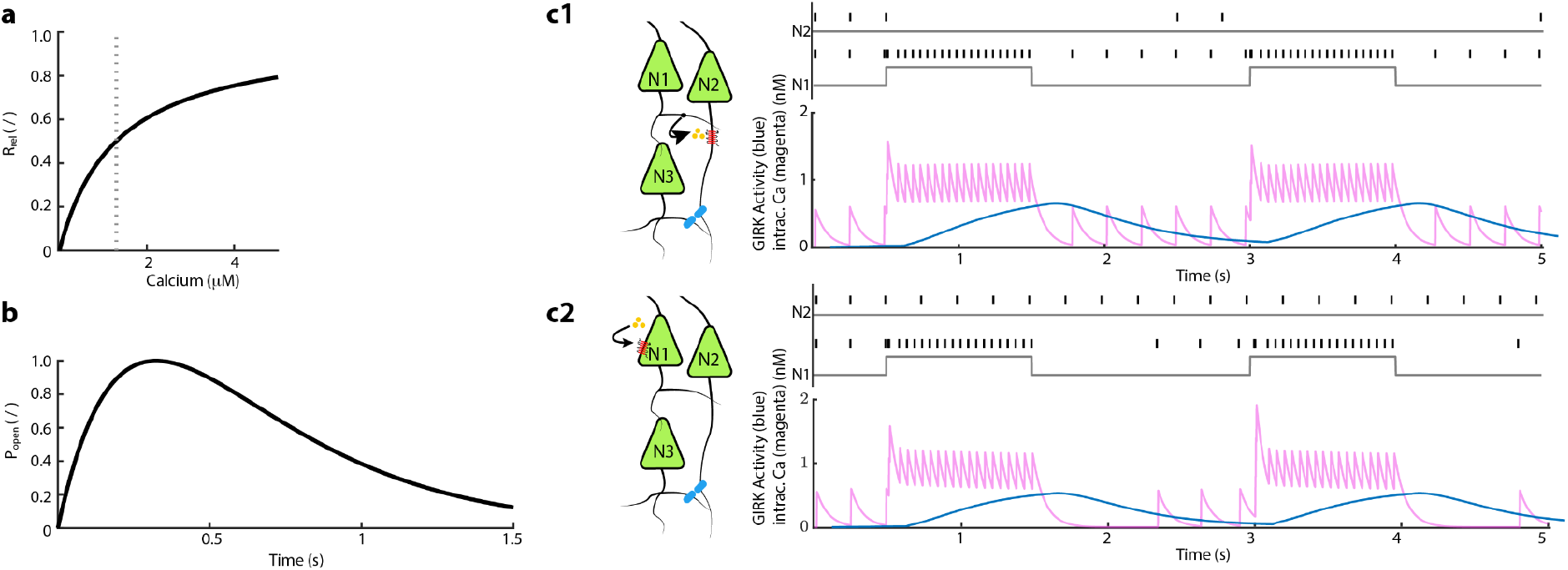
Simulation of spike rate from two LC neurons with a built-in NE-dependent GIRK activity leads to a reduced spiking rate in LC neurons that are connected via dendro-somatic synapses or a somatic release that retroact onto the secreting LC neuron. **(a)** R_rel_ is the release fraction that is dependent on the level of presynaptic calcium at the time of action potential. The gray vertical line is the amount of calcium accumulated in a cell when firing at 20Hz. The relation is defined by equation 7. **(b)** P_open_ shows the effective time course of the opening and closing of the synaptic conductance of GIRK channel. (equation 8 P _open_(t)). **(c1)** Depicts a simulation scenario where neuron 1 (N1) connects to neuron 2 (N2) via dendro-somatic synaptic interaction. *The upper diagram* shows the spike rates of N1 and N2 throughout the simulated 5-sec. Here, the *gray line* represents the time course of current injections (1s, I= 270nA) in the respective neuron. *The lower diagram* shows the development of intracellular calcium levels in N1 *(magenta*) during a phasic-like activation; the *blue line* represents the time course of GIRK conductance in N2. (**c2**) Cartoon illustrates the autoinhibition scenario in which alpha2-adrenergic autoreceptors in N1 activate through local NE-release from the soma. One-second bouts of current injections (I=270nA) lead to an increased spike rate in N1, while N2 displays a spontaneous firing pattern *(upper diagram). The lower diagram* shows time courses of intracellular calcium concentration and GIRK conductance in N1.

Based on these assumptions, we first tested our model for a synaptic-like dendro-somatic interaction between two LC neurons. To mimic episodes of high activity, we injected current amplitudes for 1-sec that lead to spike rates close to 20Hz in neuron 1 (N1) (figure 4 c1). As expected, an increased spike rate led to an uprise of intracellular calcium levels (figure 4 c1 *magenta trace* and supplemental figure 5 a,b). As described in equation 5, this increase translates into a delayed, slowly developing onset of GIRK conductance equation 10, (figure 4 b, c1 *blue trace*). The generated GIRK conductance decreases with Tau = 350ms after the current injection step ends. In our model, the increased GIRK conductance during and after episodes of high spike activity led to complete inhibition of spontaneous spiking in the dendro-somatically connected neuron 2 (N2). Only after GIRK conductance returns to basal levels, N2 resumes spontaneous spiking.

In a different scenario, we investigated how the somatic release of NE impacts the neuronal activity of the releasing neuron (N1) itself. Similar to our previous constraints, we induced phasic-like spiking through a current-step injection in our modeled neuron. The high spike rate induces an increase of intracellular calcium followed by a delayed onset GIRK conductance (figure 4 c2). Also, here, we assumed quantal release concentration as in the previous scenario, i.e., an initial extracellular NE concentration of 200nM. Despite the slowly rising of GIRK conductance in N1, the calcium concentration in N1 only shows a subtle decline in the average intracellular calcium levels during these current-induced bouts of high neuronal activity *(compare 4 c magenta traces upper versus lower trace*). The spike rate in N1 remains stable despite rising potassium leak through GIRK channels. Yet, we observed that after the current step subsided, the slow-decaying GIRK conductance impede spontaneous firing in N1; 500ms after the current pulse ended, spontaneous spiking recurred. Such an observation agrees with the typical quiescent period of LC neurons after episodes of high activity (Williams et al. 1984).

We then began to examine if somatic NE release and the spatial confinements of the LC architecture can give rise to volume-mediated interaction between LC neurons. Here, we hypothesize that the LC neurons during episodes of high activity can modulate spontaneous spiking of neighboring neurons via bystander-inhibition. The spatiotemporal NE concentration profile originating from a point source can be estimated through equation 12 (Cragg and Rice 2004; Rice and Cragg 2008).

The attenuation factor (*C*_0_ / *C*(*r*,*t*)) relative to the initial NE concentration shows a spatiotemporal profile with an attenuation of 0.001 for distances of around 20 to 25μm after 200 to 800ms (figure 5 a). The fraction of NE bound alpha2-receptors (K_D_ = 1.25nM) is shown as a temporal profile for various distances in figure 5 b. The onset and fractions of activated alpha2-receptor are slowed down and decreased through the time course of diffusion, respectively. In our neural model of LC neurons, we aim to estimate the neighbor inhibition of freely diffusing NE onto a neuron (N3) at a distance of 25μm. Evoking a phasic-like activity in neuron N1 through repeated current-step injections evoked an uprise of intracellular calcium level that surpasses the threshold for NE vesicle release. The onset of a slow-rising GIRK conductance in N3 is delayed by 1400 ms compared to previous scenarios (dendro-somatic and autoinhibition) (compare figure 4 c1, c2 versus figure 5 c). The decelerated rise of GIRK conductance is due to the small number of active alpha2-receptors. This translates into a slow and small increase of GIRK conductance in N3. As GIRK activation was delayed, a second current step two seconds after the first one ended, continued to build up GIRK conductance on the already existing conductance from the previous high activity bout (figure 5 c). Shorter intervals between the current step further facilitates this effect. This cumulated increase in GIRK conductance in N3 leads to an enlarged interspike interval between spontaneous spikes in N3 (figure 5 c and supplemental figure 6 a).

**Figure 5.**
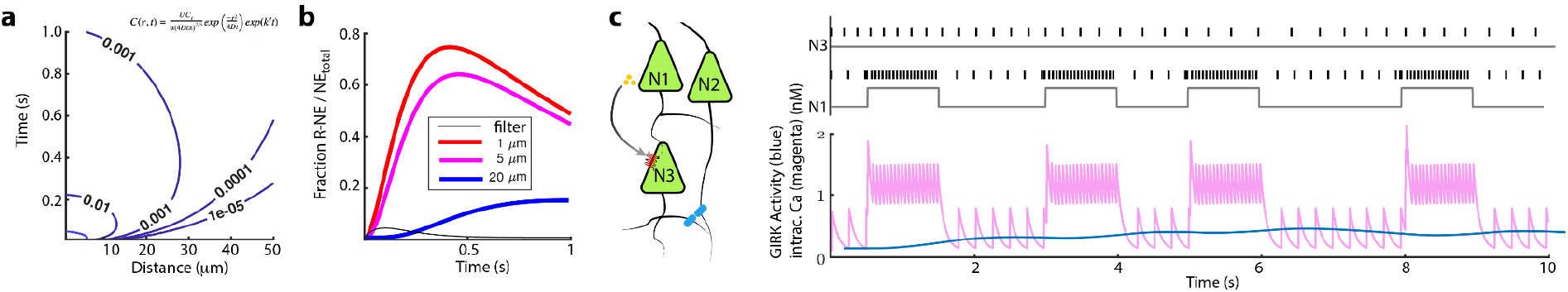
Spike modulation in the neighboring neuron of somatically release NE cumulatively affects neighboring neurons within a 25μm radius. **(a)** Spatiotemporal diffusion profiles of NE from a point source. The diffusion constant for NE was 3.4 × 10^-6^ cm^2^s^-1^ (Rice and Cragg 2008; Cragg and Rice 2004). **(b)** Time course of fractions of NE bound alpha2-receptor at 1, 5 and 20μm distance from release site. Kinetics of transmitter binding and release was modeled by an alpha function (think black line, *τ*=50 ms). **(c)** Simulation of spike train of current injected neurons (N1) and nearby neurons (N3 / 25μm distance). The lower plot shows the development of intracellular calcium levels in N1, while the blue line represents GIRK conductance in N3.

**Figure 6:**
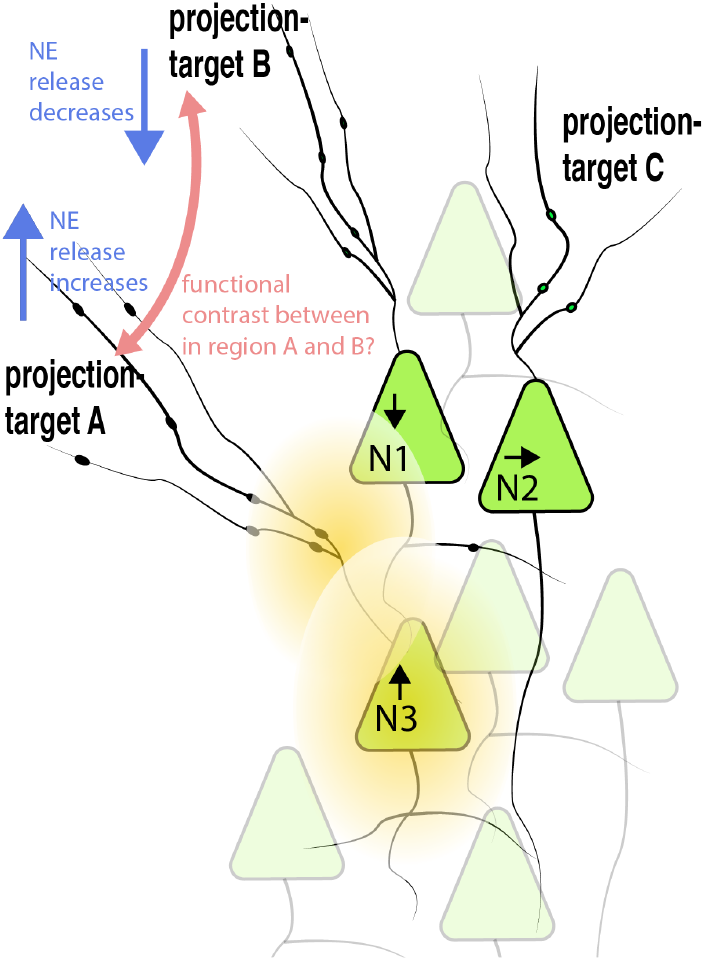
Illustration of functional consequences of a volume-mediated inhibition of neighboring neurons with different projection targets. Strong enduring input to neuron 3, N3, leads to an increase in neuronal spiking (*arrow up*) and subsequently increase release of NE within the core of the LC. Based on distances and receptor portfolio of neuron 1 (N1) from the release sites of N3, the spontaneous spiking activity in N1 will be decrease through activation of GIRK (arrow down); while for example activity in neuron 2 (N2) is not altered (arrow horizontal). Consequently, axonal terminals from N3 to projection-target area A increase spike-dependent release of NE, while axons in area B from N1 decrease NE release due to decrease spiking in N1. It is conceivable that the volume-mediated local NE inhibition in the core of the LC, leads to contrasting functional effects in the neuronal circuits in region A and B i.e. facilitating sensory salicieny in region A versus region B.

## Discussion

Noradrenergic modulation exerts robust control over neuronal circuits throughout the entire brain. The densely packed core of the LC integrates various inputs from cortical, subcortical, and brain stem areas. These inputs are computed within the nucleus and evoke various network states in the LC. Early work highlights the importance of electronic coupling between noradrenergic neurons that can synchronize sub-threshold membrane fluctuation and spiking rates in the entire LC network (Christie and Jelinek 1993; Christie, Williams, and North 1989; Ishimatsu and Williams 1996; Alvarez et al. 2002). While later research points to multiple asynchronous neuronal ensembles that dynamically switch spiking patterns within the LC (Uematsu et al. 2017; Hirschberg et al. 2017; Totah et al. 2018). As the specific neuronal architecture and synaptic wiring onto and between LC neurons are unknown, the space of potential network states and their respective output function remains elusive. Recently, the identification of local inhibitory neurons that control the phasic activity of specific neuronal subsets in the LC adds another layer of computational capacity to this brain stem nucleus (Breton-Provencher and Sur 2019; Jin et al. 2016; Kuo et al. 2020). Yet, to achieve a deeper understanding of the functional role of noradrenergic subnetworks, we need experimental approaches that target these ensembles.

Here, we developed an experimentally validated neuronal model of noradrenergic cells that allowed us to investigate NE-mediated modulatory effects between LC neurons. First, our immunostaining against DbH and TH enzymes confirms the expression of both key enzymes for synthesize of NE in the vast majority of LC neurons in young and old mice (figure 1). As recent studies demonstrated that noradrenergic axons in the dorsal hippocampus predominantly release dopamine (Kempadoo et al. 2016; Takeuchi et al. 2016), our results demonstrate that in somatic and dendritic compartments the NE can be produced in all LC neurons. Still, such partition between dopamine and NE release site could appear on the level of axonal varicosities; either through targeting of DbH enzymes to distinct projection areas or a differential expression of monoamine transporters that bias uptake of a specific catecholamine from the extracellular space.

Our reconstruction of LC geometry reproduces noradrenergic cell numbers that are five times lower than earlier histological studies (Sturrock and Rao 1985;O’Neil et al. 2007). As previous studies use stereological cell counts of 6 to 50μm brain slices, overestimation through double counting of neurons is possible. Yet, it is likely that these large differences are also due to insufficient penetration of TH antibodies in our cleared tissue preparation that limit cell count compared to classical histology. Nevertheless, a possible underestimation of the cell count in our LC preparation would only affect the overall fraction of LC neurons that are in a 25μm-radius, but the resolved absolute number of LC neuron pairs would stay constant or be larger. In our diffusion model, we have not incorporated noradrenaline transporters activity (NET) that can limit the diffusion radius of NE. Despite studies showing the presence of transporters in the LC core (Klimek et al. 1997; Isaias et al. 2011), previous diffusion modeling studies for the Substantia nigra argue that the impact of monoamine transporter on the diffusion radius of dopamine is small (Cragg and Rice 2004; Rice and Cragg 2008). Direct electrophysiological recordings in the LC neurons confirm that increasing electrical evoked-release of NE scales with the rate of alpha2-receptor evoked inhibitory potentials in noradrenergic neurons; arguing that monoamine transporters are not drawing excessive NE out of the extracellular space (Courtney and Ford 2014). In contrast, in similar dose measurements in the ventral tegmental area, D1-receptor evoked inhibitory responses that did not show dependence on stimulation strength. Only after treatment with a DAT antagonist responses are scaling with stimulation strength (Courtney and Ford 2014). These experimental findings argue that NET activity within the LC is not limiting the diffusion of NE.

The observed reduction in spontaneous spike rate in our model through the diffusion of extracellular NE has so far not been described to our knowledge; it has been discussed as a plausible mechanism for dopaminergic neurons in the substantia nigra as well as dopaminergic axons (Cragg and Rice 2004; Rice and Cragg 2008; Walters et al. 2014). Key factors for the observed bystander-inhibition are distances and geometrical orientation of the release site between LC neurons. Our model deduces that extracellular NE can accumulate through episodes of long-lasting phasic activity or repeated periods of bursting discharge. This accumulation is dependent on the speed of Gi signaling cascade, GIRK channel kinetic and dispersion of extracellular NE through the densely packed LC core. Such bystander-inhibition adds another layer of interaction between LC neurons. The physiological role of such type of inhibition is strongly interwoven with a topographical organization of the LC.

The idea of topographic organization of the LC has been put forward decades ago; Most studies at that time used local injection of retrograde dyes (Waterhouse et al. 1993; Loughlin, Foote, and Bloom 1986; Loughlin, Foote, and Fallon 1982). Despite controversial findings, it is generally accepted that the LC has a dorsal-ventral organization in which neurons in the dorsal part of the LC are preferential projecting to frontal areas while the ventral part projects to brain stem areas. For an example, LC neurons that project to hippocampal areas or olfactory bulb are more likely found in the dorsal part, while ventral LC neurons are more probable to project to medulla or spinal cord (Mason and Fibiger 1979; Loughlin, Foote, and Bloom 1986; Schwarz et al. 2015). Based on new viral strategies and high-throughput axonal mapping technologies, a better understanding of the brain-wide input-output organization of the LC became available. These findings helped to develop new concepts that now allow also to understand the controversial findings from earlier tracing studies. For example, one of these concepts argues that axonal projections from a single LC neuron forms an extensive axonal field to a specific brain area, while also possessing collaterals to other brain areas at lower density. A variation of such a concept entails the distinction between broadcasting LC neurons that form equally dense innervation to many areas, versus exclusive LC neurons that form specific innervation to a single brain area. Both concepts would blur retrograde tracing experiments, and underestimate a topographical organization of the LC. Interestingly, functional studies highlight heterogeneity of projection-defined LC neurons in terms of intrinsic excitability, after-hyperpolarization and their proteomic make-up (Wagner-Altendorf, Fischer, and Roeper 2019; Chandler, Gao, and Waterhouse 2014). Here, LC neurons that preferentially project to the prefrontal cortex or hippocampus show a different sensitivity to alpha2-receptor agonists. Therefore, a topographic organization and differential expression of adrenergic receptors could fine-tune the impact of neigboring inhibition. It is interesting to speculate how the here postulated volume-mediated interaction of spatially clustered LC neurons with different projection targets could amplify differences in extracellular NE in their respective axonal fields. The increase in contrast of NE levels in two target regions could increase sensory saliency in one region.

## Data availability

All code used for all analyses and plots are publicly available on GitHub at https://github.com/teamprigge

## Acknowledgments

We are thankful to the Center for Brain and Behavioral Science for financing K.M. M.P is supported by the Best Minds programs from the Leibniz Association. We are in depth to Menachem Segal for discussion and Oliver Kobler from the Combinatorial NeuroImaging (CNI) core for support with light sheet and confocal microscopy.

## Author contributions

SB, HH, KM, TF, JB and MP conceived experiment(s) and developed the analyses. MP wrote first draft of manuscript with support SB, KM and JB. MP and JB supervised the work, and acquired funding.

## Supplement

**Supp. Figure 1:**
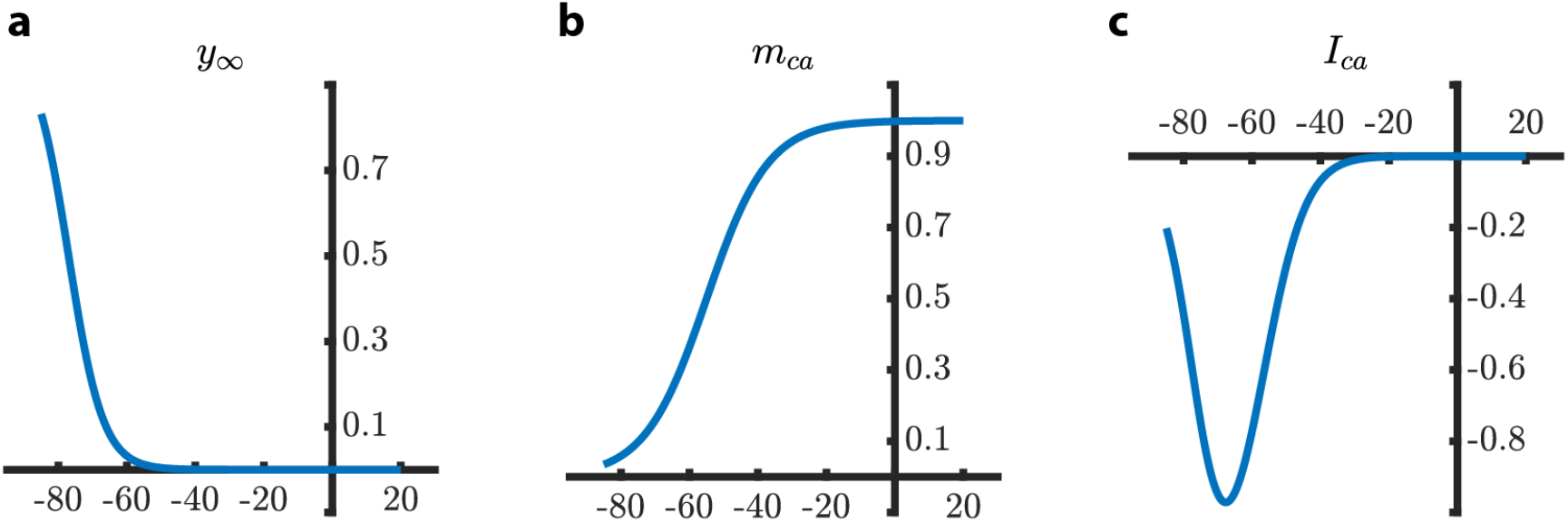
The calcium current is responsible for the spontaneous spiking of the cell. It is unclear in Alvarez et al if this is a transient conductance (with activation and inactivation gates) or a persistent conductance (with two kinds of activation gates). If we take the formulas as given, The ‘fast’ gating variable mCa opens with membrane depolarization. Specifically, it approaches unity for membrane potentials above −40 mV (see Supp Figure 1). The ‘slow’ gating variable y∞ closes with membrane depolarization. Specifically, it approaches zero for membrane potentials above −60 mV. In the same regime, the characteristic time to approaches 20 ms. Taken together, this current is active in a window between −80mV and −40mV, being strongest around −60mV. As a result, this current can cause a slow Ca-spike, on which multiple fast Na-spikes can occur. Activation variables for y∞ (left) and m_ca (middle) and I_ca (right) for calcium current, as a function of membrane voltage, for the formulas as given by Alvarez et al.

**Supp. Figure 2:**
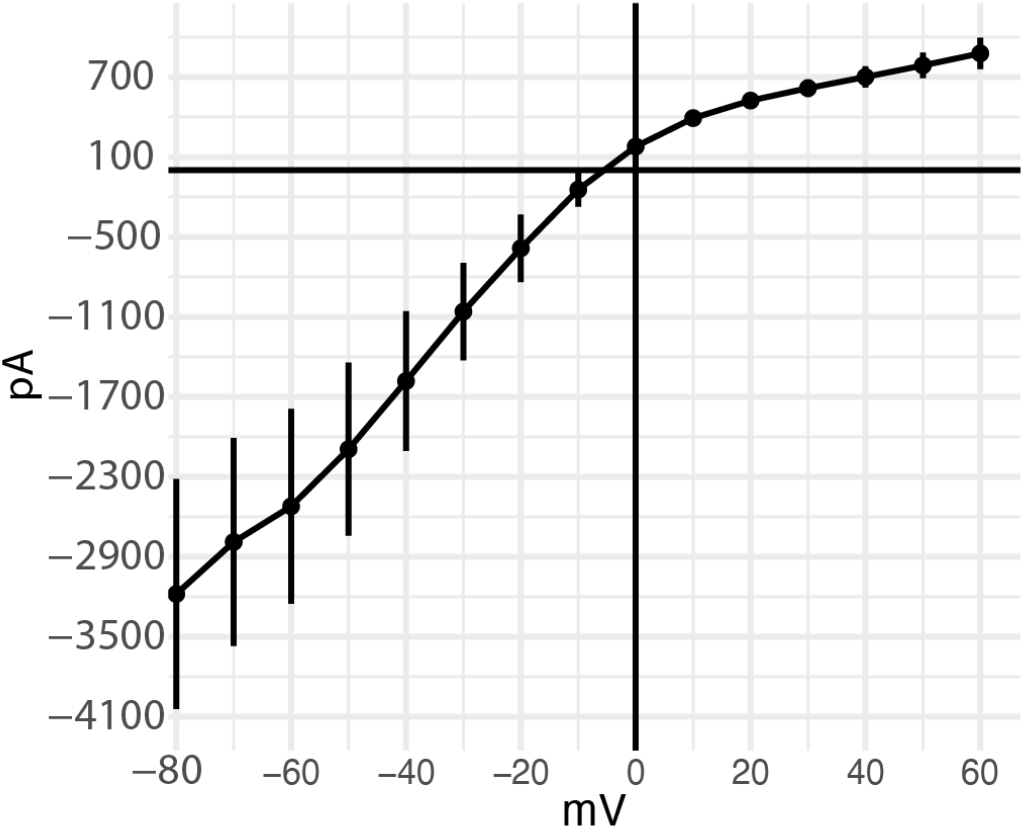
Inward-rectification presented in a current-voltage plot derived from HEK293 that heterologus express GIRK2 channels. GIRK time course are used in equation 13.

**Supp. Figure 3:**
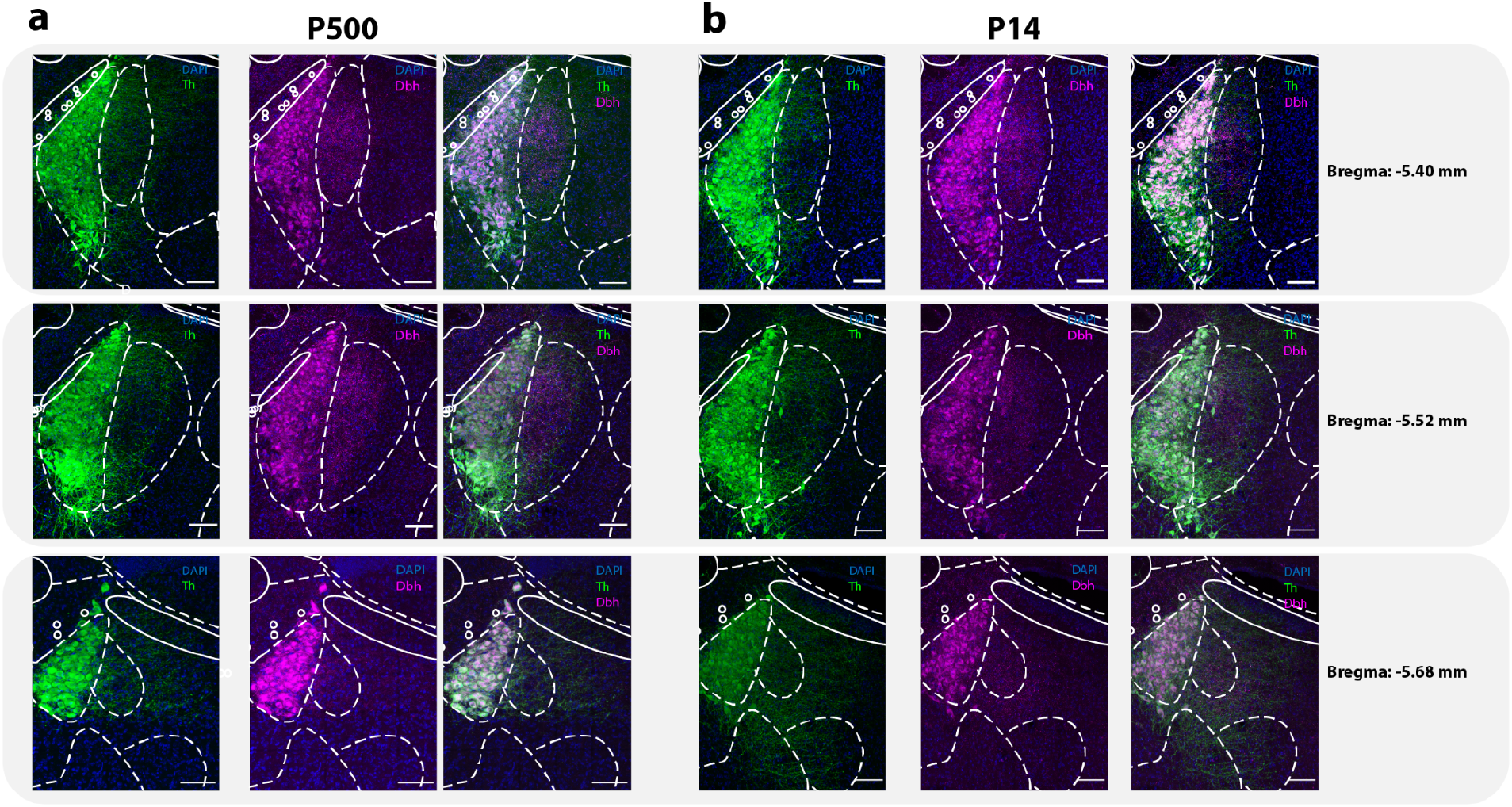
Confocal images of immunostained brainstem slices against tyrosine hydroxylase (TH / green) and dopamin-beta-hydroxylase (DbH / magenta) at three different position from bregma: −5.40, −5.52 and −5.68mm. **(a)** Shows LCs slices from an old animal (P500), while **(b)** overviews LC slices from a young animal (P14). Somatic as well as dendritic compartments exhibit different expression levels for TH and DbH within the LC topography.

**Supp. Figure 4:**
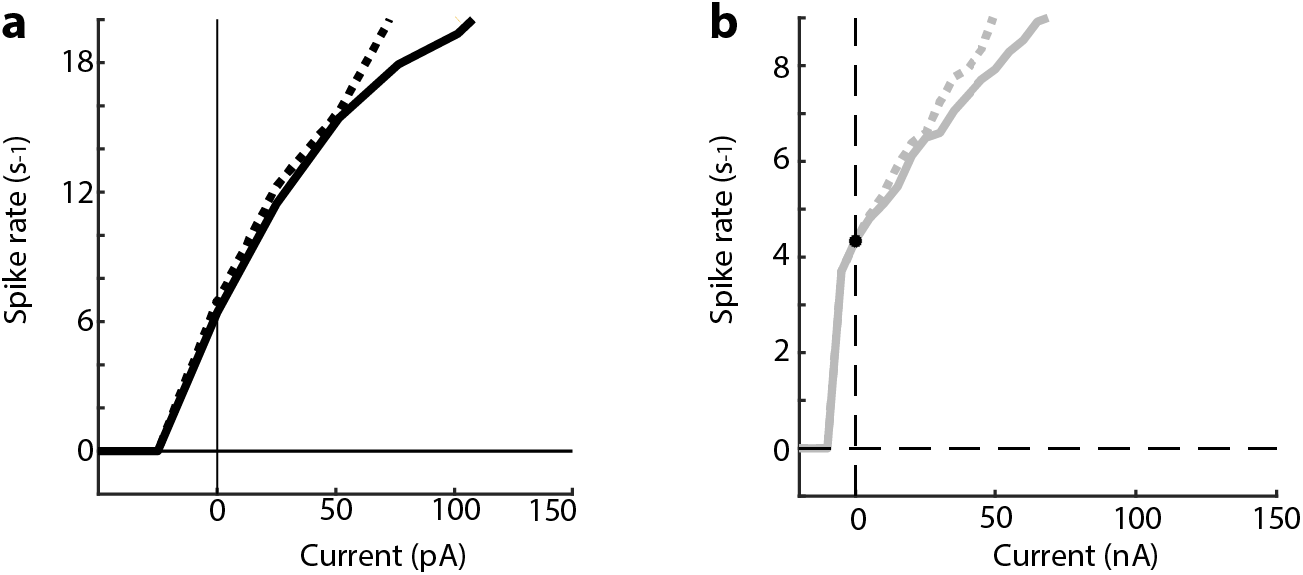
Spike rate dependence on 300-ms current-injection steps separated for early spikes (spike 1 and 2 / dotted line) versus later spikes (> spike 2 / full line). **(a)** FI Curve of an experimental neuron. It is firing at almost 6.1Hz at spontaneous state. **(b)** FI curve of a simulated neuron. It is firing at almost 4.5Hz at spontaneous state.

**Supp. Figure 5:**
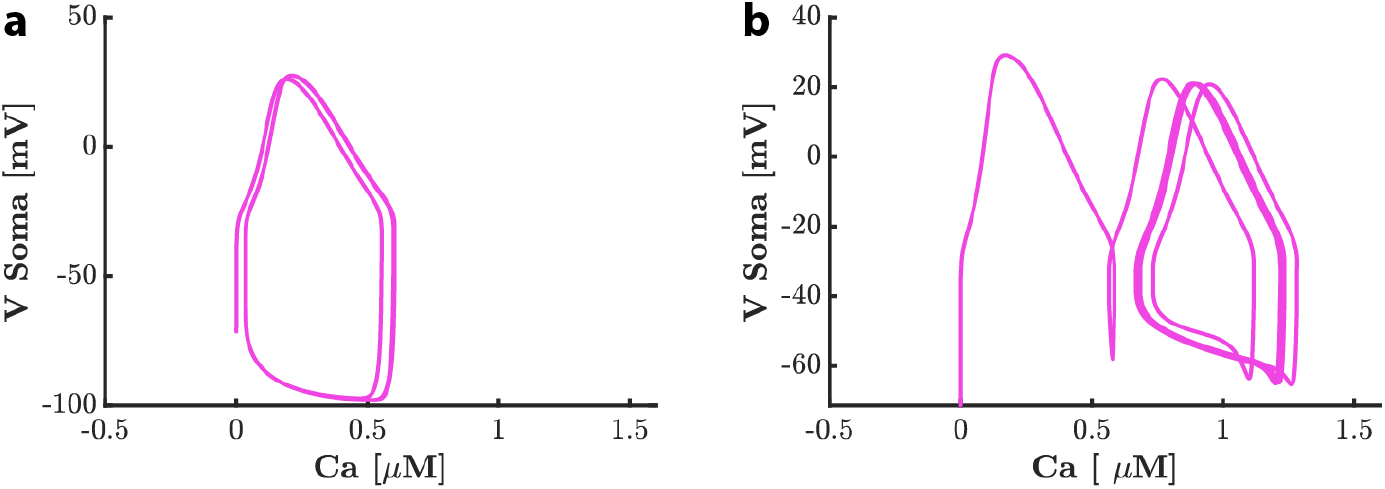
Action potential triggers calcium accumulation in the cell which facilitates the release of neurotransmitters along with many other calcium-triggered events. When the simulated neuron is firing at 4Hz, a small amount of calcium is accumulated and equibrilates at constant level as shown in fig 3 **(a)**. When the same neuron is firing at 20Hz, a higher level of calcium is accumulated after several action potentials **(b)**.

**Supp. Figure 6:**
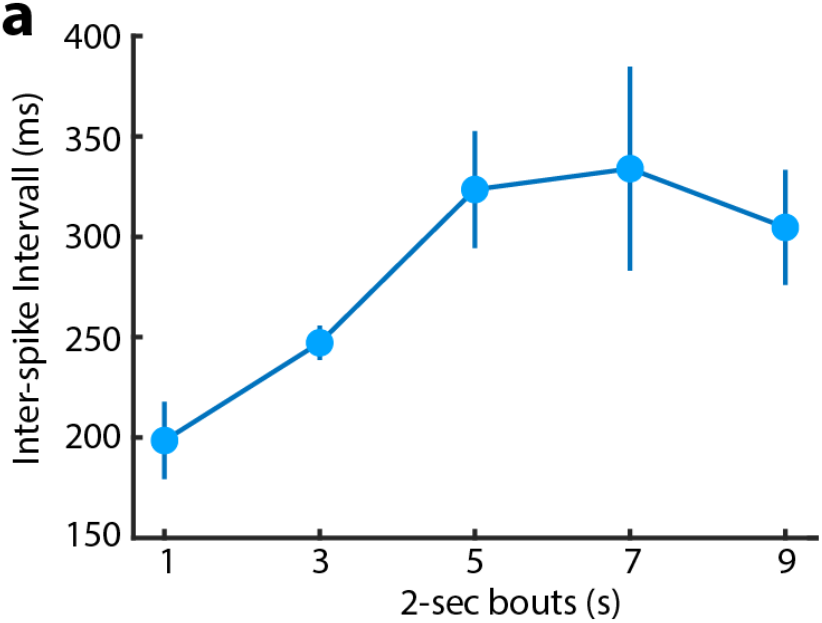
Effect of extracellular NE that diffuses freely in the extracellular space. The released NE binds to a receptor site in a nearby neuron and triggers opening of GIRK. Unlike synaptic inhibition, inhibition by diffusion is delayed in nature as it takes time to reach the binding site when diffused in extracellular space. Here ISI (binned 2s) are calculated for neuron N2 in figure 5. Inhibition slowly builds up (1s-7s) after N1 releases NE during several intervals which leads to a significantly longer ISI in N3. N3 slowly gets back to the steady state after N1 stops releasing NE(9s).

